# Preservation of soluble enhanced green fluorescent protein (EGFP) within fixed NIH 3T3 fibroblasts

**DOI:** 10.1101/2025.09.22.677869

**Authors:** John T. Elliott, Mahir Mohiuddin, Alex Tona

**Affiliations:** Cell Systems Science Group, Biosystems and Biomaterials Division, National Institute of Standards and Technology, Gaithersburg MD 20899

**Author notes:** Corresponding author John T. Elliott.

**Keywords:** Fixative, MBS, DSG, DSP, PFA, GFP, NIH 3T3, Jurkat, image analysis, automated microscopy, fluorescence, enhanced green fluorescent protein, EGFP, quantitative measurements

## Abstract

Quantitative imaging of cytoplasmic green fluorescence protein (GFP) in fixed cells can be unreliable if the fixing process does not preserve the total fluorescence intensity level or the spatial location relative to the living cells. In this study, we examine the effect of fixatives (formaldehyde, disuccinimidyl glutarate (DSG), dithiobis(succinimidyl) propionate (DSP) and m-maleimidobenzoyl-N-hydroxysuccinimidyl ester (MBS)), fixative buffers and the cross-linking times on the fluorescent intensity of soluble enhanced GFP (EGFP) expressed within NIH 3T3 cells. The total fluorescence intensity within individual cells during the fixation process was measured in an automated fluorescence microscope. Our results show that choice of fixative, the fixing solution and the cross-linking time were important for minimizing EGFP losses during fixing. The optimal fixation condition for these cells was identified to be the MBS cross-linker in a microtubule stabilizing buffer. After an 8 h fixation, greater than 90 % of the initial GFP fluorescence within cells was preserved. This was 3-fold higher than the GFP fluorescence remaining when the cells were fixed with 1 % paraformaldehyde in PBS. MBS treated cells could be permeabilized with 0.05 % Triton X-100 with little additional loss in fluorescence intensity. The MBS fixative could also preserve soluble fluorescent proteins in suspension cells, suggesting the fixation results are not cell line-specific. Direct imaging of the fixation process on fluorescent reporter cells provided insight into the effect of chemical fixatives on biological cells and can facilitate identification of fixation protocols that are fit-for-purpose for quantitative measurements.

## Introduction

Fixation procedures that preserve the cell physical state are important for many microscopy investigations in cell biology. It is important that the desired structural features of the fixed cell and the location and concentration of the specific molecules being examined are representative of those in the living cell[1]. Most fixatives used in microscopy experiments function as non-specific protein cross-linkers[2]. The cross-linking is thought to covalently connect cellular proteins and stabilize cytoskeletal components. A typical fixative for cell culture is paraformaldehyde (PFA) in phosphate buffered saline (PBS)[3, 4]. Although it functions well in many applications, several comparative studies have indicated that some fixatives may outperform others when preserving specific cellular structures and intracellular components[1, 5, 6]. Fixing conditions should be optimized for a specific application to assure data can be collected on representative samples during cellular imaging.

The introduction of fluorescent protein (FP) technologies has greatly facilitated the measurement of molecular activity within cell[7]. For example, FPs are often expressed as fusion protein to study protein location or expressed as soluble cytoplasmic FP that report on gene promoter activity[8, 9]. In these reporter cells, the total FP intensity can be measured in individual single cells and used to describe the distribution of promoter expression within a population of cells. This measurement is valuable for studying phenotypic and epigenetic heterogeneity in cellular populations with tools such as high content imaging.

Measuring intracellular FP concentrations in fixed cells will depend on how well the fixative preserves the FP content in the live cell[10]. Although paraformaldehyde and organic solvent-based fixatives are used extensively to fix cells with FP, they have been observed to significantly reduce the fluorescence signal associated with cells expressing a cytoplasmic soluble FP[6, 11]. Directly evaluating the effect of fixation on the intracellular FP fluorescence intensity is valuable for establishing optimal fixing conditions that enable quantitative fluorescence measurement relationship between corresponding live and fixed reporter cells[1].

In this study, we examined the use of fixatives on a clonal population of the TN1-pd2EGFP-1/NIH 3T3 reporter fibroblast cell line that expresses a cytoplasmic destabilized enhanced green fluorescent protein (EGFP) in response to activation of the Tenascin-C promoter[8]. These cells have an increase in total cellular fluorescence when they are actively proliferating and exhibit a wide range of fluorescence intensities to provide a sufficient distribution of intensities. The effect of the fixing process on the fluorescence intensity measured from these cells was quantitatively monitored *in situ* to identify an optimal fixing condition.

## Methods and Materials

### Fixatives and Buffers

Phosphate buffered saline (PBS) and microtubule stabilizing buffer (MTSB) were used as fixing solutions. PBS was obtained from Thermo Fisher Gibco. MTSB[5] is a mass fraction of 4 % polyethylene glycol (PEG) 8000, 100 mM 1,4-piperazinediethanesulfonic acid (PIPES), 10 mM ethylene glycol-bis(2-aminoethylether)-N,N,N’,N’-tetraacetic acid (EGTA), pH 6.9. Paraformaldehyde solution (16 % in H_2_O) was obtained from Electron Microscopy Sciences (Hatfield, PA). The bifunctional crosslinkers disuccinimidyl gluterate (DSG), dithiobis(succinimidyl) propionate (DSP) and m-maleimidobenzoyl-N-hydroxysuccinimidyl ester (MBS) were obtained from Thermo Fisher Pierce and MilliporeSigma. The cross-linkers were dissolved in cell grade DMSO and added to either the MTSB or PBS fixing buffers immediately before adding to the cells. The stock and working concentrations used for NIH 3T3 cells are described in Table 1.

**Table 1.**
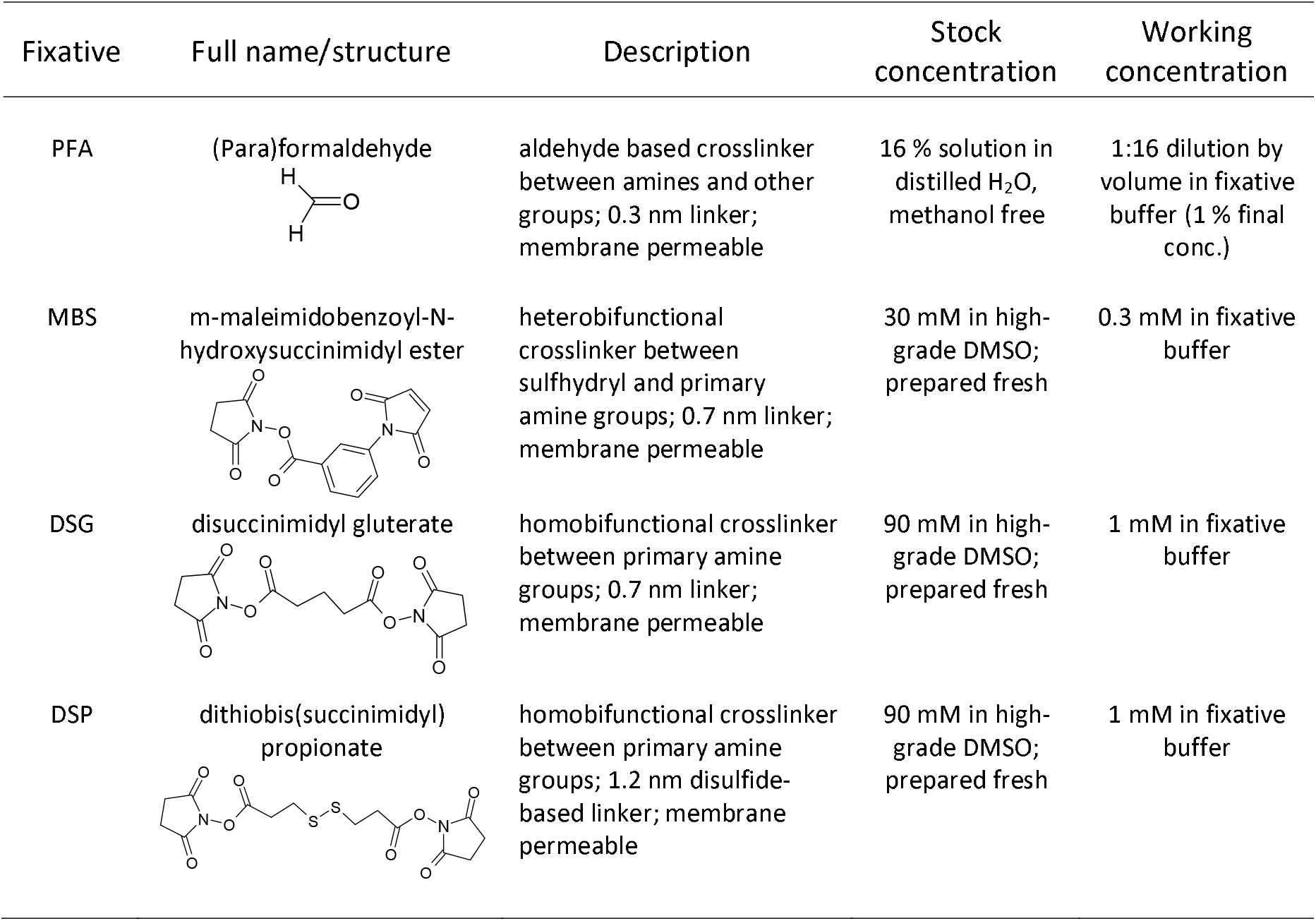
Fixatives used in this Study.

| Fixative | Full name/structure | Description | Stock concentration | Working concentration |
| --- | --- | --- | --- | --- |
| PFA | (Para)formaldehyde | aldehyde based crosslinker between amines and other groups; 0.3 nm linker; membrane permeable | 16 % solution in distilled H<sub>2</sub>O, methanol free | 1:16 dilution by volume in fixative buffer (1 % final conc.) |
| MBS | m-maleimidobenzoyl-N-hydroxysuccinimide ester | heterobifunctional crosslinker between sulfhydryl and primary amine groups; 0.7 nm linker; membrane permeable | 30 mM in high-grade DMSO; prepared fresh | 0.3 mM in fixative buffer |
| DSG | disuccinimidyl glutarate | homobifunctional crosslinker between primary amine groups; 0.7 nm linker; membrane permeable | 90 mM in high-grade DMSO; prepared fresh | 1 mM in fixative buffer |
| DSP | dithiobis(succinimidyl) propionate | homobifunctional crosslinker between primary amine groups; 1.2 nm disulfide-based linker; membrane permeable | 90 mM in high-grade DMSO; prepared fresh | 1 mM in fixative buffer |

### Cell and Sample Preparation

TN1-pd2EGFP/NIH 3T3 reporter cells were generated and maintained as previously described[8]. Before cell seeding, the 10 cm TCPS dishes (BD Biosciences) were inverted and two 3 cm strips of transparent office tape were placed perpendicular to each other near the corner of the plate. The inside corner of the crossed tape pieces served as a fiduciary mark for resetting the stage position and one of the pieces of tape served to align the plate to the automated stage. Cells were seeded on the dishes at 800 per cm^2^ and allowed to adhere overnight. The low seeding density facilitates measuring the total integrated intensity within the cells with an image analysis software.

### Imaging the Fixation Process

The cells were fixed within 2 d to 3 d of seeding. A single 10 cm dish was removed from the incubator and fitted into the stage of a Zeiss Axiovert 100 inverted microscope with phase and fluorescence capabilities. The fluorescence was excited with a 100 W Hg lamp and imaged with a Photometrics CoolSnapFX CCD camera though a dichroic filter set optimized for EGFP fluorescence. An automated stage allowed repeated movement to a specific location on the culture dish relative to a fiduciary mark. The stage was moved to a field with 10 cells to 20 cells, and the spatial coordinates of the stage were stored. The culture media was directly removed by aspiration while the TCPS plate was in the microscope stage. The cells were rinsed with 20 mL of Hepes-Hanks buffer (Thermo Fisher Gibco) at 37 °C. Phase contrast and fluorescence images of the cells were collected and used to measure the fluorescence signal levels in the live cells. The Hepes-Hanks buffer was removed by aspiration before the fixative was added. Phase and fluorescence images were typically collected from the cells before and after every step in the fixing procedure using the instrument settings identical to those used for initial imaging of the live cells. If several cell culture dishes were being imaged for replicate and comparison experiment, the dish was removed from the stage and the next dish was aligned and imaged.

After various timepoints (1 h, 3 h or 8 h) of fixation at room temperature (RT), a dish was placed on the microscope stage and moved to the location of the stored coordinates. Phase and fluorescence images were collected. The fixative was removed and 0.05 % Triton X-100 (TX-100) in fixing buffer was added to the cells. Images were collected after 30 min. The detergent solution was removed and the cells were rinsed twice with PBS (pH 7.4) plus 0.05 % sodium azide. Phase and fluorescence images were again collected. In some cases, cells were fixed with a single fixative (MBS) for > 8 h and then treated with a second fixative (PFA) before imaging.

### Quantifying Intracellular GFP

All image analysis was performed with ImageJ-FIJI software[12]. The fluorescence and phase images that were compared were aligned manually using the ImageJ software plugin TurboReg[13]. In general, cells were segmented using the phase image and the regions of interest were transferred to the corresponding fluorescence images. Areas adjacent to the cell that did not contain cell fluorescence were used to determine the average local background intensity. The mean integrated GFP intensity within each cell was calculated by first subtracting the average local background intensity from each pixel in the cell area and then averaging the adjusted intensity of the pixels in the cell area. The fraction of the original mean GFP fluorescence intensity remaining within the cells was determined by plotting the before-fixation mean intensity verses the after-fixation mean intensity on a cell-by-cell basis and fitting the data with a linear regression (Microsoft Excel). All quantitative experiments were performed with at least 3 replicates prepared in separate petri dishes. Each replicate included images of 10 cells to 100 cells for calculating changes in intracellular fluorescence intensity.

### Microscope Quality Control and Other Images

The total excitation power of the incident lamp on the microscope was measured by placing a lightmeter probe (Newport) directly over the objective lens with the EGFP dichroic filter in place. The power output measured was typically 6.9 mW for the 100W Hg lamp. Although lamp intensities were stable for hours, random fluctuations (up to 5 %) were observed due to the lamp source. We also imaged photostable fluorescent Schott Glass filters (GG475) to ensure that the fluorescence intensity measured at the CCD camera did not change significantly day-to-day during the experiments[14]. A flatfield control was not used for this study as the measured fluorescence intensity over the imaging area did not vary more than 5 % due to incident illumination alignment.

Higher magnification images of fixed cells exhibiting extracellular vesicles (i.e. blebs) or intracellular structures were collected with a 40X objective. The f-actin cytoskeletal structures of the fixed cells were imaged by incubating the cells in 1:200 dilution (volume fractions) Texas Red-labeled phalloidin (Thermo Fisher) in 0.05 % TX-100 in PBS. The cells were rinsed with PBS and imaged through a filter cube optimized for the Texas-Red dye.

The quenching effect of PFA on the GFP fluorophore (see supporting information S1) was determined by quantitative imaging of the GFP fluorescence signal from MBS-MTSB fixed reporter cells before after the addition of up to 1 % PFA volume fraction in PBS. Quantitative fluorescence images of the cells after exchange of the PFA solution to a PBS solution were collected to calculate the GFP fluorescence signal recovery after removal of the PFA fixative. The fluorescent signals in the images were quantified using ImageJ as described above.

The photobleaching rate of the GFP signal in MBS-MTSB fixed cells was determined by continuously exposing the fixed cells to incident fluorescent light from the objective for up to 3 min. Fluorescent images of the cells were collected every 15 s and the fluorescent signals were quantified using ImageJ as described above (see supporting information S2).

### Fixation of GFP-containing suspension cells

A fluorescent Jurkat cell line (VCN-4) was maintained as described previously[15]. Cells (20 × 10^6^) were harvested into a 15 mL conical tube, rinsed with PBS, suspended in MBS-MTSB fixative (0.6 mM MBS in 10 mL MTSB) and placed on a tube rotator (10 rpm) overnight. Alternatively, cells were suspended in PBS containing 4 % volume fraction PFA. After fixation, cells were pelleted (300 *g*, 5 min) and rinsed three times with PBS containing 1 % mass fraction bovine serum albumin (PBS/1 % BSA). Fixed cells (2 × 10^6^ /mL) were stored in PBS/1 % BSA at 4 °C for at least 2 weeks. Live VCN-4 cells were pelleted, rinsed and resuspended in PBS/1 % BSA at the same concentration. Fluorescent cells (50 000) before and after fixation were compared using a flow cytometer (BD Accuri) with no additional sample preparation. At least 3 replicate experiments were performed to estimate loss in GFP fluorescence after cell fixation.

## Results

A list of the fixatives evaluated in this study are shown in Table 1. These cross-linkers were chosen because they are cell membrane permeable and exhibit reactivity with functional groups available on most proteins. Cross-linkers with the N-hydroxysuccinimide groups and maleimide groups react more selectively with amine and sulfhydryl groups, respectively[2]. Formaldehyde under aqueous conditions can cross-link amines and other functional groups[16]. The cross-linkers in this study were dispersed in phosphate-buffered saline (PBS) or a microtubule-stabilizing (MTSB) buffer.

### Quantifying the Retention of Intracellular GFP after Fixation

The NIH 3T3 reporter cells used in this study express a soluble cytoplasmic EGFP when the tenascin-C promoter is activated. Figure 1a shows a plot of the total integrated intensity of individual cells in a single field of view before and after fixation with several of the buffer and cross-linker combinations used in this study. The slope of the fitted line is the average fractional amount of initial GFP fluorescence remaining in the cells after fixation.

**Figure 1.**
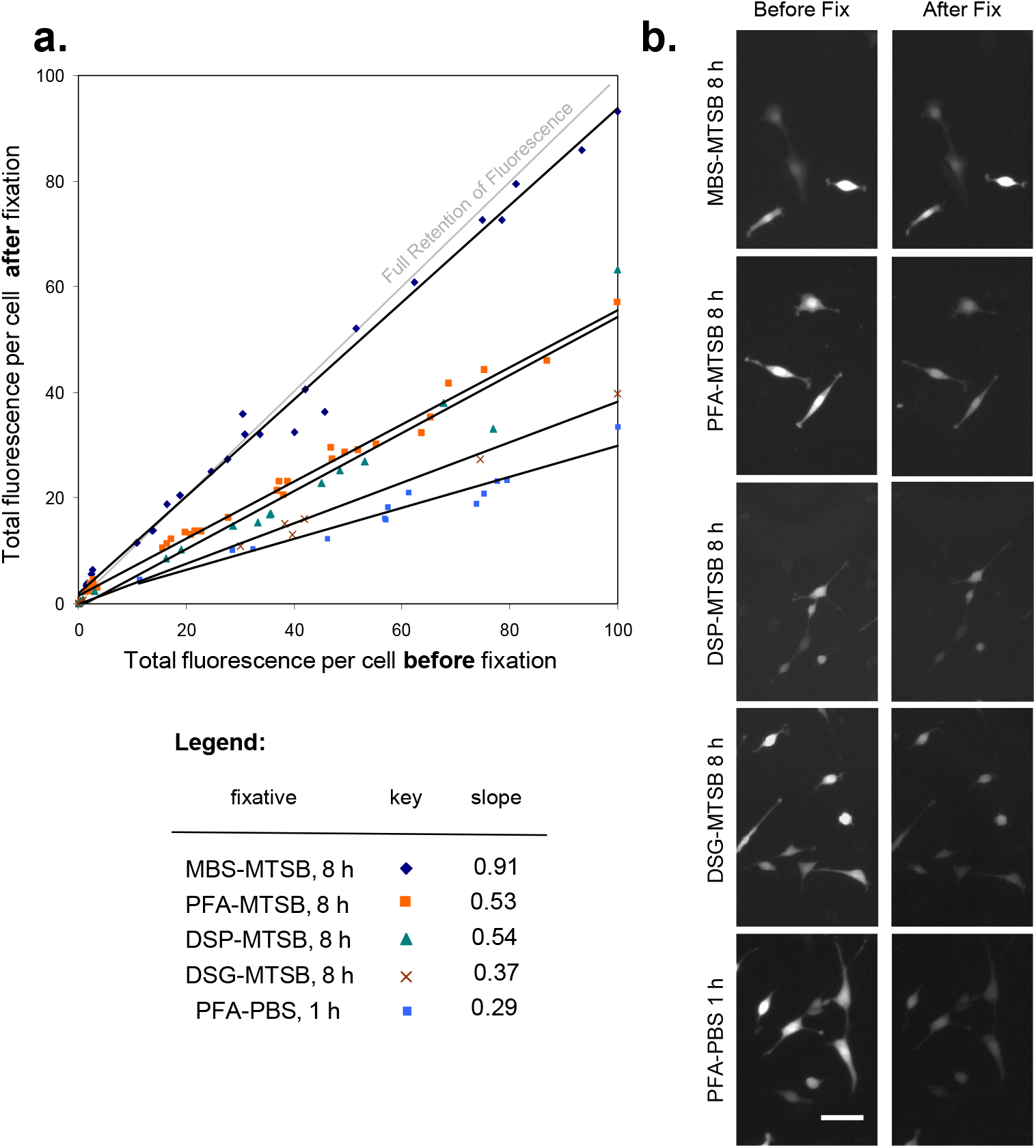
The effect of fixation on total GFP intensity measured within cells. A. Representative data plotted as total fluorescence within cells before and after fixation. The slope correspond to the lines fitted through the data (all regression coefficients greater than 90 %. B. Representative images of GFP containing cells fixed with the fixatives shown in the legend. The white bar is 80 µm.

A fixation procedure commonly used for cell culture studies is addition of 1 % PFA in PBS for 1 h at RT[17]. As can be seen in Figure 1a, this fixation condition results in a 71 % reduction in the observed GFP fluorescence. This was the least optimal fixation condition identified in this study. When cells were treated with 1 % PFA in MTSB for at least 8 h, the fractional remaining fluorescence was increased to 52 %. Fixation with the amine-sensitive bifunctional cross-linkers disuccinimidyl gluterate (DSG) and dithiobis(succinimidyl) propionate (DSP) in MTSB preserved 37 % and 54 % of the GFP in the cells, respectively. An interesting fixing condition was found when the MBS fixative was used with MTSB buffer. The plot in Figure 1a shows that the measured GFP intensity within cells after fixation is greater than 90 % of the pre-fixation intensity.

Each of the fixation conditions examined in this study showed that there is a high correlation between the individual cell intensities before and after fixation (R^2^> 0.95). This linear relationship indicates the relative intensity of GFP between each of the living cells in the population is preserved during fixation despite significant losses in the average cell intensities. Figure 1b shows representative images of GFP fluorescence in cells before and after fixation.

A summary of the effects from different fixatives, fixative buffers and fixation times on the preservation of GFP fluorescence is shown in Figure 2. The use of MTSB instead of PBS as a fixing buffer during an 8 h fixation improved the preserved GFP intensity for all fixatives. The effectiveness of the cross-linker in MTSB decreased in the following order: MBS > PFA = DSP > DSG. Extending the fixing process from 1 to 8 h significantly improved the preserved GFP intensity for most fixing procedures. Further increasing the fixing time to 16 h did not further improve the GFP preservation (data not shown). The addition of a second fixation treatment (1 % PFA-MTSB for 1 h at RT) to cells that were previously fixed with MBS-MTSB for 8 h did not significantly influence the intensity of recovered fluorescence.

**Figure 2.**
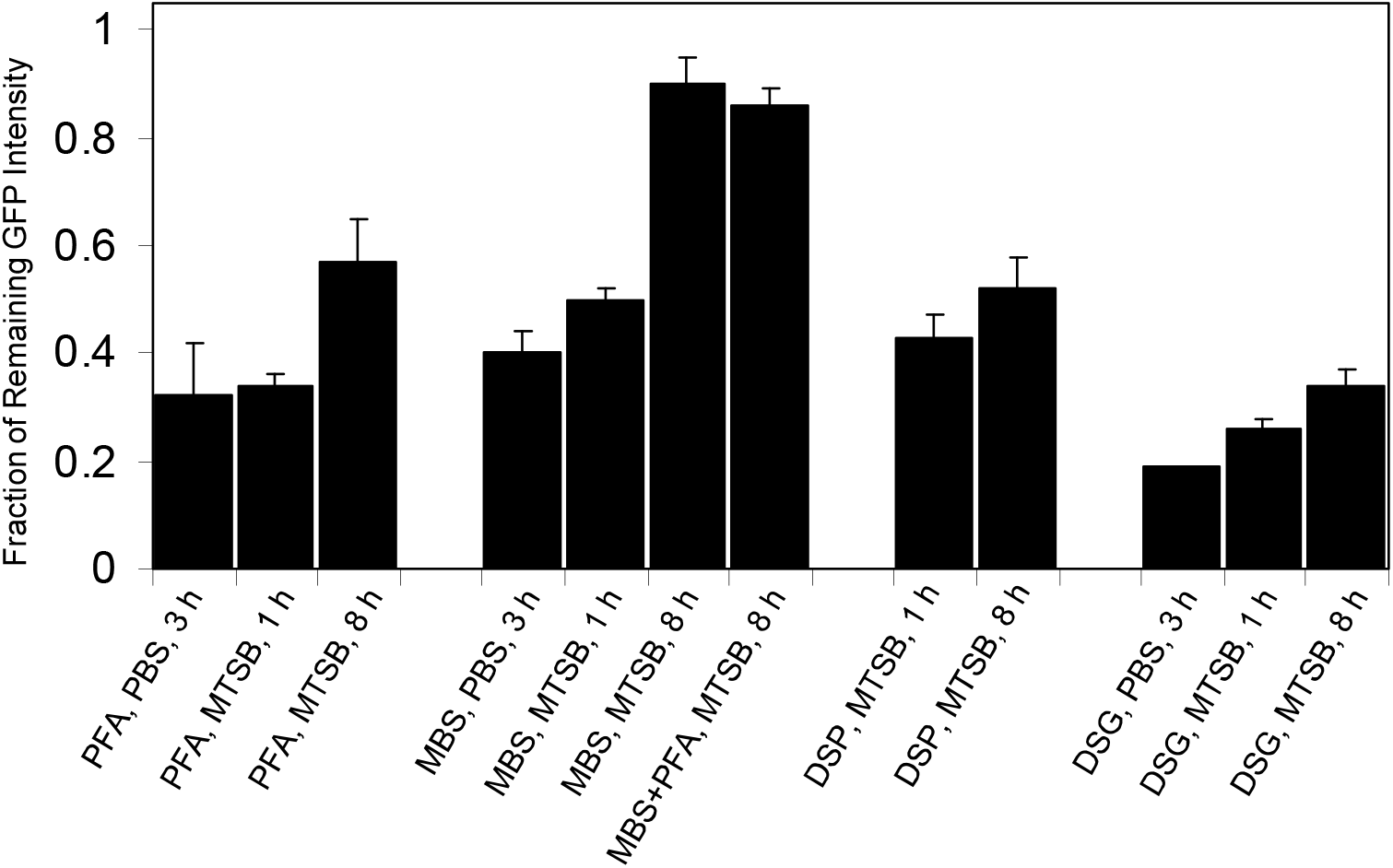
Summary of average fixation conditions on the fraction of GFP fluorescence preserved within NIH 3T3 cells. Each SD error bar represents the variation from separate measurements from 3 different plates.

Identifying Losses in Cytoplasmic GFP Intensity during Fixation: Figure 3 shows the measured fluorescence intensity within cells as a function of time during the fixation steps with several conditions. The fluorescence intensity within the cells is reduced by 10 % and 27 % within 5 min after the addition of the MBS and DSG fixatives in MTSB, but the intensity drops 40 % after the addition of PFA in MTSB (time point 1). An additional 3 % to 5 % of fluorescence is lost during the 8 h fixing time (time point 2). When the cells are rinsed and permeabilized with Triton X-100 (TX-100), the DSG treated cells lose an additional 15 % of fluorescence intensity, where the PFA and MBS fixed cells show only an additional 5 % loss in fluorescence (time point 3). When TX-100 is rinsed out from the cells, the PFA and MBS treated cells show an increase in fluorescence (time point 4). The DSG fixed cells show only a small change in fluorescence intensity after removal of the detergent. A final rinse to remove the MTSB fixing buffer and replace it with Hepes-Hanks results in an additional 5 % to 15 % increase in fluorescence for each of the treatments (time point 5). This study indicates that the changes in GFP fluorescence occurs at several steps during the fixation process, but the largest decrease in fluorescence is dependent on the fixative.

**Figure 3.**
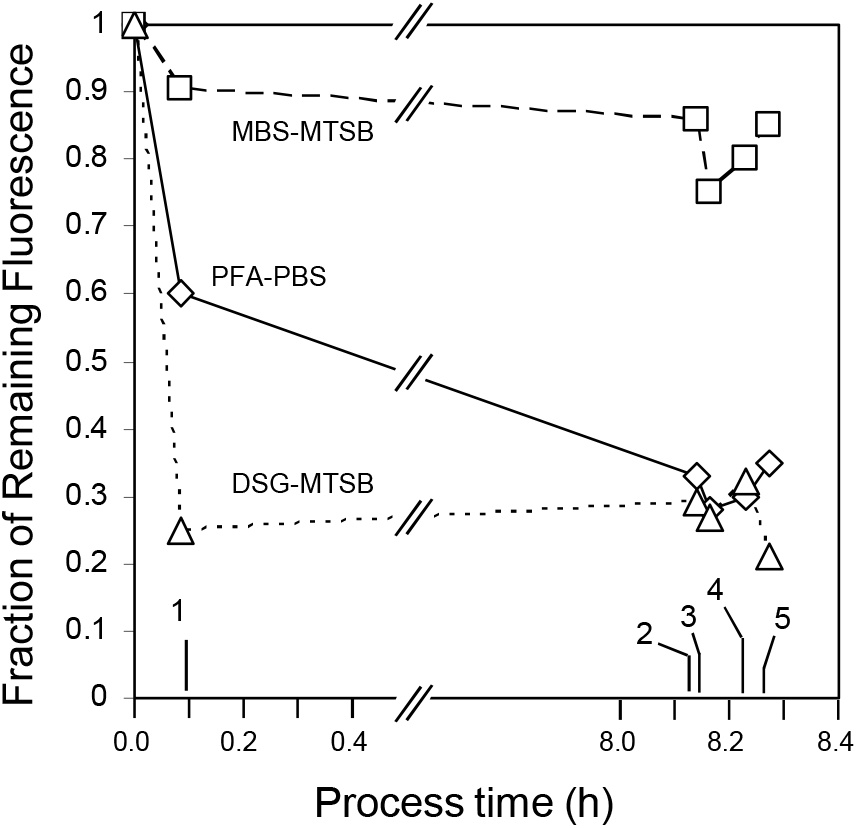
The effect of the fixation process on the total GFP intensity measured within NIH 3T3 cells. Each plot is from a representative data set. 1) 5 min after addition of fixative, 2) 8 h after addition of fixative, 3) addition of TX-100 in MTSB buffer, 4) rinse with MTSB buffer, 5) rinse with HEPES Hanks. Values at each point are referenced to fluorescence intensity values measured for living cells in HEPES Hanks buffer before fixation. Under some fixing conditions, bleb formation on cells would be observed at time point 1.

A possible mechanism that may reduce fluorescence from soluble GFP in fixed cells is the formation and release of extracellular vesicles (i.e. blebs) filled with cytoplasmic protein[16]. Figure 4 shows phase contrast images of blebs formed on the cell surface during the DSG treatment. When the same cells were imaged by fluorescence microscopy, the vesicles contain a significant amount of soluble GFP protein. Some of the vesicles became several microns in size and in some cases, they detached from the cells. Upon treatment with TX-100 detergent, the vesicles lysed immediately. Blebs were also observed within 10 min after the addition of DSG and within 30 min after the addition PFA and DSP in MTSB. Vesicular structures were not observed with cells treated with MBS-MTSB.

**Figure 4.**
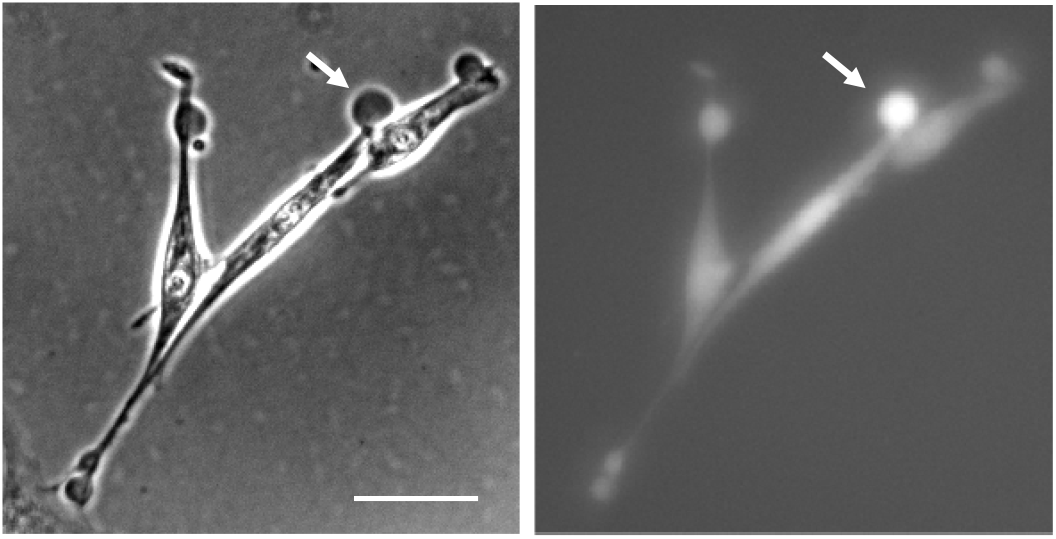
Loss of soluble EGFP during the formation of extracellular vesicles (i.e. blebs). Some fixing procedures resulted in the formation of vesicles on the cell surfaces (white arrow). Some of these vesicles became several μm in diameter and fluorescence images revealed they contained soluble cytoplasmic GFP. These vesicles disappeared after buffer exchanges and TX-100 treatment suggesting a loss of intracellular soluble GFP during fixing maybe due to bleb formation. White scale bar is 10 µm.

We also examined if the chemical fixative itself can influence the measured GFP fluorescence intensity. When fluorescent cells were first fixed with MBS-MTSB and then treated with any of the other fixatives in PBS, only the PFA fixative caused a significant decrease in fluorescence (≈ 19 % decrease with 1 % PFA, see supporting information S1). When the PFA was removed from the cells during rinsing, complete recovery of the GFP fluorescence was observed. This result indicates that the presence of PFA can reversibly quench the GFP fluorophore, but it does not cause permanent chemical damage of the GFP molecule.

It is important to note that the preserved GFP from any fixation processes was highly photostable. Less than 4 % of the initial intensity was lost after continuous exposure of the cells to excitation light for 2 min (Supporting Information S2). In this study, the cells were typically exposed for less than 3 s during a measurement.

### Qualitative Morphology and Cytoskeletal Structure Evaluation

All of the fixative conditions generally preserved the overall shape and spread morphology characteristics of the cells (Figure 1b). Figure 5 shows high-magnification images of cells fixed with MBS-MTSB and PFA-MTSB and stained with phalloidin. As seen, the location of the GFP within the cytoplasm and the intracellular appearance of the F-actin stress fibrils are similar for cells treated with either fixative.

**Figure 5.**
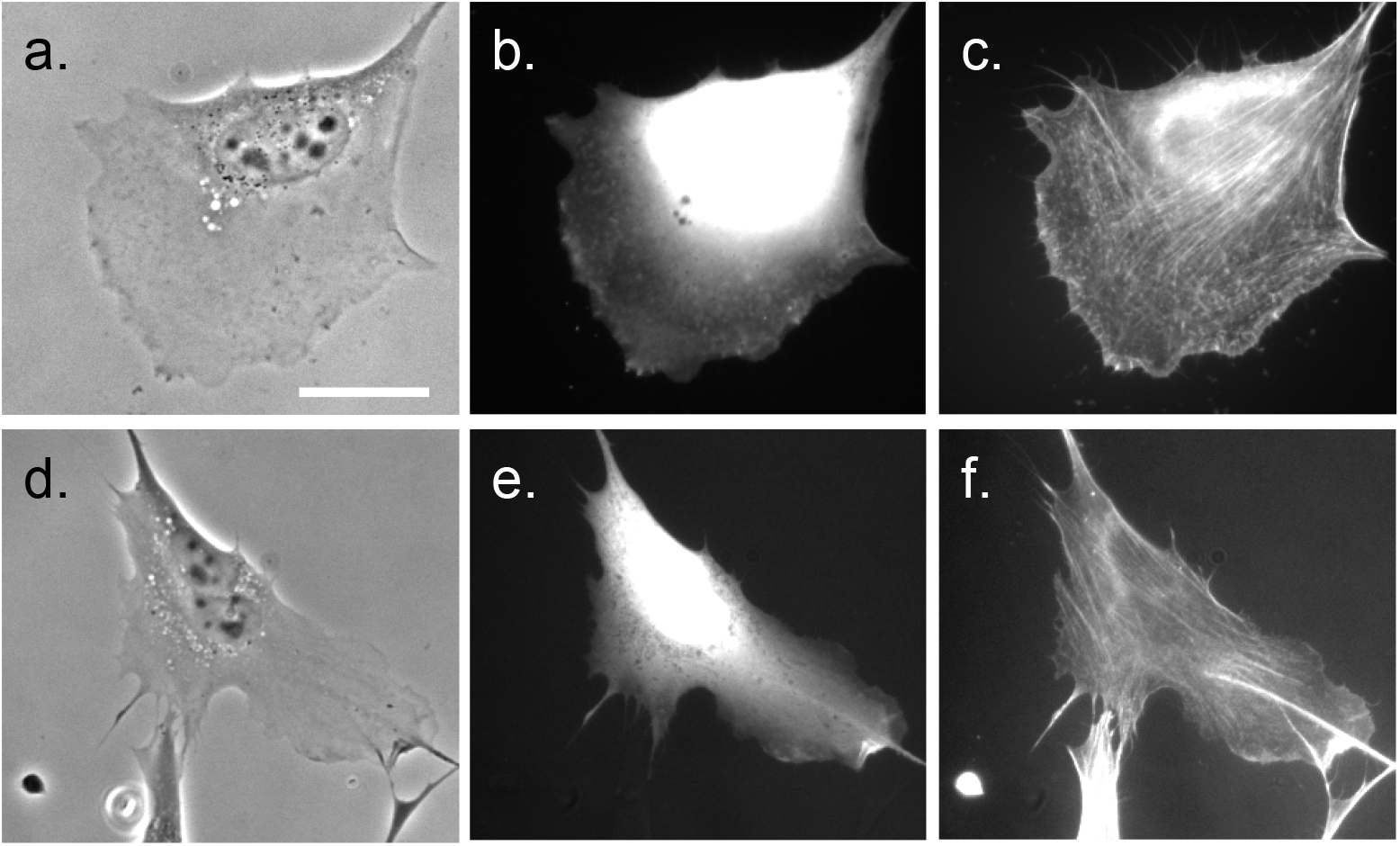
NIH 3T3-TN1dEGFP cells were fixed overnight with 1 mM MBS (a-c) or 1 % PFA (d-f) in MTSB and stained with Tx-Red phalloidin. Panel a and d are phase images, panel b and e are images of the cytoplasmic GFP fluorescence. Panels c and f are images of phalloidin staining. Thin f-actin stress fibers are observed in cells under both fixation conditions. White scale bar is 10 μm.

### Fixation of other GFP-containing cells

The MBS fixation process also preserves intracellular GFP and morphology in a suspension cell phenotype. A 10 % loss in GFP signal and similar shape-dependent scattering profiles were observed before and after MBS-MTSB fixation of fluorescent Jurkat cells (VCN-4, Supplemental Figure 3)[15]. Fixation of the same cells with PFA in PBS resulted in greater than 60 % loss in fluorescence and appears to alter the scattering profile of the cells. These results are consistent with the fixation effects observed in the fluorescent NIH 3T3 cells (e.g. Figure 2) and suggests that MBS-MTSB fixative may preserve intracellular soluble GFP in other cell types.

## Discussion

Development of preservation methods that stabilize biological structures within cells is important for cell observations and assays that require robustness with respect to sample handling and storage time[10]. Although fixatives such as PFA are used as general cell preservative, a variety of fixatives and fixing conditions have been shown to differentially stabilize biological structures such as transcription factors, cytoskeleton, integrins and intracellular features[1, 5, 18]. The NIH 3T3 reporter cell line used here contains an engineered gene that expresses a non-fused and destabilized EGFP during activation of the tenascin-C promoter[8]. EGFP is a fluorescent protein that has been enhanced to improve fluorescence properties and monomer solubility[19]. The expressed protein is a 28 kDa β-barrel approximating a 400 nm x 25 nm cylinder structure. For this study, the FP was assumed to be freely soluble in the cytoplasm with a homogeneous concentration across the cell. An ideal fixative for performing measurements on EGFP fluorescence would preserve the concentration of the FP β-barrel within the cell and not affect the fluorescence signal.

By using *in situ* fluorescence microscopy to directly monitor the fixing of reporter cells, we were able to quantitatively assess the effectiveness of each fixation procedure. A 70 % loss in intracellular EGFP fluorescence was observed when 1 % PFA in PBS was used as a fixative. Interestingly, this loss was reduced to 40 % with 1 % PFA in the MTSB buffer. This is consistent with the idea that the fixing buffer plays a functional role in the preservation process[5]. We also tested more selective crosslinkers that conjugate molecules through specific functional groups (Table 1). Although cell treatment with the homobifunctional amine-crosslinkers DSP and DGS has been shown to stabilize cytoskeletal and other biological structures[20, 21], their use in this study resulted in a greater than 30 % loss in intracellular EGFP fluorescence. Surprisingly, MBS, which cross-links free primary amines to free sulfhydryl groups, was able to preserve greater than 90 % of the intracellular FP signal when used in the MTSB buffer. This directly highlights the ability to tailor fixation conditions to best preserve cellular features of interest.

To identify the possible origins of fluorescence loss during fixing, control experiments were performed to test for chemical changes in the EGFP fluorophore due to photobleaching or quenching and visible structural changes in the cells. EGFP was photostable and was not irreversibly quenched with the chemical fixatives used in this study (Supplemental Figures 1 and 2). Although general cell morphology appeared to be preserved for all fixatives (i.e. Figure 1), direct imaging of the fixation process revealed that extracellular blebs are formed on the cells in the presence of most fixatives (i.e. Figure 4). Fluorescence images show that the micron-size blebs contain cytoplasmic EGFP, and they disappear by the end of the fixing process.

We hypothesize the much of the fixation-induced decreases in cellular fluorescence is due to the loss of GFP in the bleb structures. Previous studies have shown that soluble cytoplasmic proteins are encapsulated in the blebbing vesicle structures generated with fixative chemicals[22, 23]. Fixation-induced blebbing likely results from cytoskeletal failures, though both the MTSB fixation buffer and the MBS cross-linker, which prevented a greater than 90 % loss in fluorescence, have been shown to affect cytoskeletal stabilization. Each of the components of the MTSB buffer (i.e. PIPES, EGTA and PEG) directly affect both cytoskeletal assembly and the osmotic flow of the cytoplasm[24]. The MBS crosslinker has been shown to specifically cross-link components of fibrillar actin and is used as a botanical fixative that stabilizes cytoskeletal ultrastructures in plant tissues[18]. It is possible that this fixative and buffer combination stabilizes the cytoskeletal structure which prevents the loss of soluble EGFP even in the presence of detergent (Figure 3). Our quantitative study indicated treatment of MBS-MTSB fixed cells with a second fixative (PFA) did not significantly change the recovered intracellular GFP fluorescence (Figure 2). This suggests that multiple fixatives could be used to preserve multiple cellular features in addition to intracellular GFP concentration. Furthermore, the MBS-MTSB fixative was also capable of preserving soluble GFP in phenotypically distinct suspension cells with a differing cytoskeletal structure. This suggests the fixation chemistry should function in a variety of cell types. The MBS-MTSB fixed NIH 3T3 cells and Jurkat cells were morphologically stable for several months and the preserved GFP fluorescence was stable for at least 3 weeks.

## Conclusion

We have identified a fixation procedure that highly preserves the concentration of soluble EGFP within adherent NIH 3T3 fibroblast cells. The method is based on the MBS cross-linker that conjugates free amine (e.g. lysine) and free sulfhydryl (e.g. cysteine) groups, in a microtubule stabilizing buffer. By directly monitoring the fixing of live FP reporter cells, mechanistic insight into how the fixation process affects the measured fluorescence intensity could be generated. Three-fold changes in the GFP fluorescence intensity within the fixed cells can be observed with different fixation conditions. These methods will be useful for fixing soluble EGFP in reporter cell lines and for understanding how fixatives can influence the preservation of biological structures for cellular measurements.

### NIST Disclaimer

Certain commercial equipment, instruments and materials are identified in this paper to specify an experimental procedure as completely as possible. In no case does the identification of particular equipment or materials manufacturer imply a recommendation or endorsement by the National Institute of Standards and Technology nor does it imply that the materials, instruments, or equipment are necessarily the best available for the purpose. This manuscript is a contribution of NIST, and therefore is not subject to copyright in the United States.

