## Supplemental Figures for "Preservation of soluble enhanced green fluorescent protein (EGFP) within fixed NIH 3T3 fibroblasts"

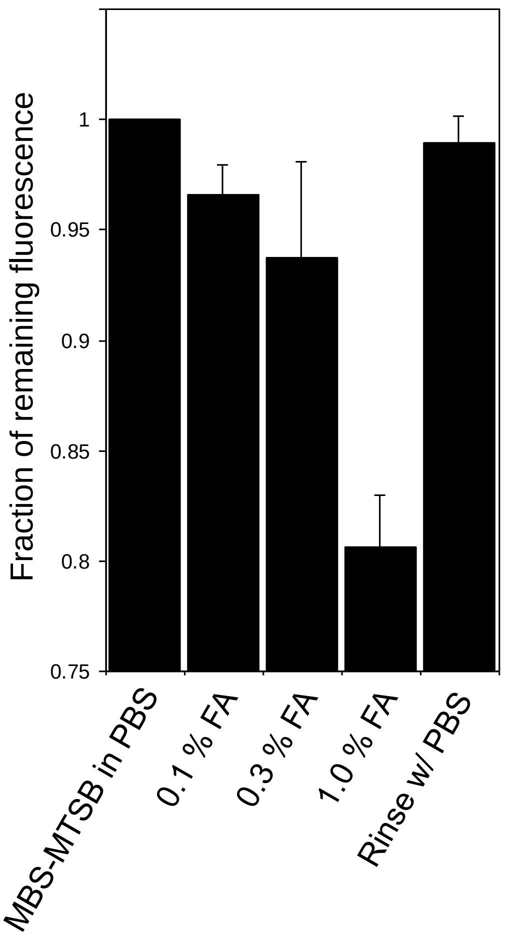


Supplemental Figure 1: MBS-MTSB fixed NIH-3T3 cells containing EGFP were treated with increasing concentrations of formaldehyde (FA). The presence of 1 % FA caused a 20 % reduction in EGFP fluorescence, but the emission was fully recovered after removing FA by rinsing in PBS. This suggests FA does not chemically modify the EGFP fluorophore but chemical quenching of EGFP fluorescence does occur in the presence of FA.

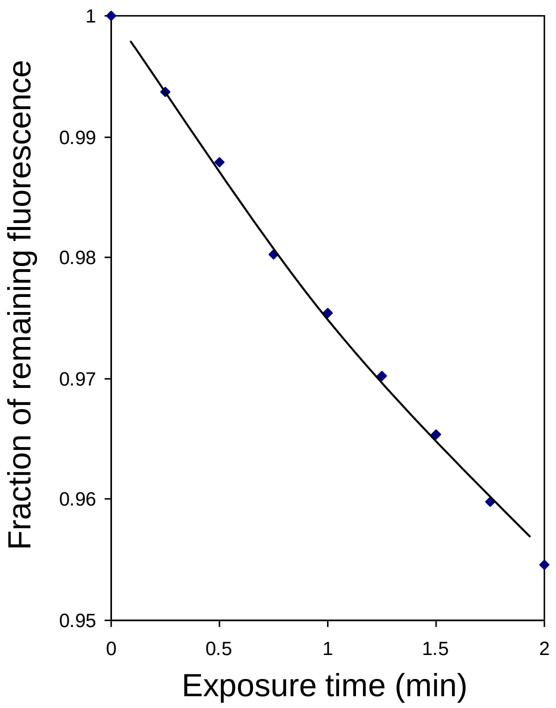


Supplemental Figure 2- Photobleaching of cytoplasmic EGFP in NIH-3T3 cells. Only 4 % of the fluorescent signal is lost after a two-minute exposure (6.2 mW). The cells in this study were exposed to incident light for less than 3 s.

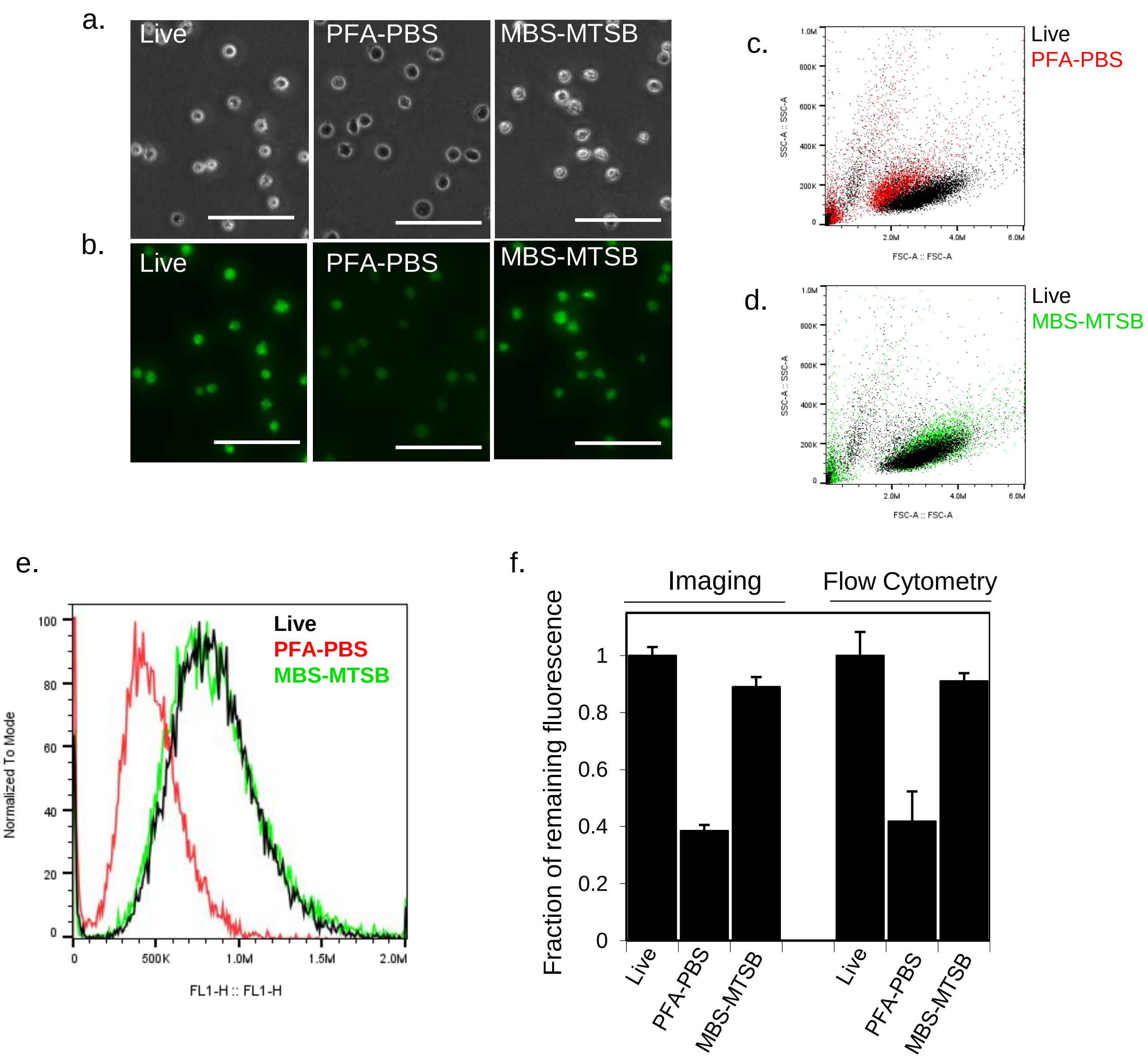


Supplemental Figure 3- A suspension-based T-cell line (Jurkat) expressing cytoplasmic soluble EGFP was fixed with PFA-PBS or MBS-MTSB. Small changes in cell morphology were detected in both phase imaging (a) and flow cytometry scattering profiles (c) of the PFA-PBS fixed cells. In addition, the PFA-PBS fixed cells exhibited a greater than 60 % loss in intracellular fluorescence as determined by quantitative fluorescence imaging (b) and flow cytometry (e, f). Fixation with MBS-MTSB resulted in minimal changes in cell morphology (a, d) and only a 10 % loss in intracellular fluorescence (b, e, f). These results are similar to those obtained with fixing adherent NIH-3T3-EGFP cells and supports the idea that the fixative could be used for many cell types expressing soluble fluorescent proteins. Scale bar = 100 µm.
